# Delineating the site of interaction on the intracellular domain of 5-HT3A receptors with the chaperone protein RIC-3

**DOI:** 10.1101/689596

**Authors:** Elham Pirayesh, Antonia G. Stuebler, Akash Pandhare, Michaela Jansen

## Abstract

The serotonin type 3A (5-HT_3A_) receptor is a homopentameric cation-selective member of the pentameric ligand-gated ion channel (pLGIC) superfamily. Members of this superfamily assemble from five subunits, each of which consists of three domains, extracellular (ECD), transmembrane (TMD), and intracellular domain (ICD). Previously, we have also demonstrated that 5-HT_3A_-ICD is required and sufficient for the interaction between 5-HT_3A_ and RIC-3. Additionally, we have shown that 5-HT_3A_-ICD fused to maltose binding protein (MBP) directly interacts with the chaperone protein resistance to inhibitors of choline esterase (RIC-3), without the involvement of other protein(s). To elucidate the molecular determinants of this interaction we developed different MBP-fused 5-HT_3A_-ICD constructs by deletion of large portions of its amino acid sequence. We have expressed seven mutants in *Escherichia coli* and purified them to homogeneity. Using a RIC-3 affinity pull-down assay, the interaction of MBP-5HT_3A_-ICD constructs and RIC-3 is investigated. Furthermore, we co-expressed 5-HT_3A_ and 5-HT_3AB_, a heteromeric form of 5-HT_3_Rs, with RIC-3 in *Xenopus* oocytes to compare their interaction with RIC-3 in-vivo by two electrode voltage clamp (TEVC) recordings. Full-length 5-HT_3A_-and 5-HT_3AB_ mediated currents are significantly reduced when RIC-3 is co-expressed in either condition. In summary, we identify a 24-amino acid long segment of the 5-HT_3A_-ICD as a molecular determinant for the interaction between the 5-HT_3A_-ICD and RIC-3.

**Statement of Significance:** The chaperone protein RIC-3 is known to modulate the functional surface expression of cation-conducting pentameric ligand-gated ion channels. Previously we have demonstrated that the intracellular domain of serotonin channels mediates this effect. Here we provide experimental evidence for a 24-amino acid long segment within the 115-amino acid long intracellular domain as a determinant for RIC-3 interaction. Recently it was found experimentally that the identified segment contains an alpha helix that has been observed or predicted to be present in other cation-conducting channels. The present work provides novel insights into protein-protein interactions that are likely also relevant for other cation-conducting members of this large ion channel family that includes nACh and 5-HT3 receptors.

## Introduction

Serotonin receptors type 3 (5-HT_3_), like the nicotinic acetylcholine (nACh), □-aminobutyric acid type A (GABA_A_), and glycine (Gly) receptors, are members of the pentameric ligand-gated ion channel (pLGIC) superfamily. Ion channels of this Cys-loop superfamily are homo- or heteropentamers. 5-HT_3_ receptors are composed of five identified subunits (A-E) and are involved in excitatory neurotransmission [1]. These receptors are abundant in the central and peripheral nervous systems (CNS, PNS) and are found pre- and postsynaptically [2]. Among all subunits of 5-HT_3_ receptors, the 5-HT_3A_ subunit is widely distributed in CNS, PNS, and internal organs and is also found in non-neural cells and tissues across the human body [3]. While the 5-HT_3B_ subunit is less abundant, the mRNA of this subunit has been detected in several regions of the human brain, intestines and in the kidney [4]. The mRNAs of the remaining 5-HT_3_ subunits (C-E) have been found in PNS and CNS, organs and extraneuronal cells [5, 6]. Currently, these receptors are mainly targeted in emesis treatment in patients undergoing chemotherapy or suffering from irritable bowel syndrome [5, 7]. Recent studies have shown potential correlations between 5-HT_3_ receptor activity and several neurological disorders such as anxiety, psychosis, nociception and cognitive function [8–10]. Some studies also suggest that 5-HT3 receptors play a role in bipolar disorder and schizophrenia [11, 12].

Each subunit of a pLGIC pentamer consists of three domains, namely extracellular domain (ECD), transmembrane domain (TMD) with four membrane spanning helices (M1-M4), and intracellular domain (ICD, only in eukaryotic pLGICs)[13]. The assembly, functional maturation and membrane trafficking of pLIGIC subunits are modulated by different chaperone proteins depending on their subtype. The protein Resistance to Inhibitors of Cholinesterase (RIC-3) is known as a modulator of nAChRs as well as 5-HT_3A_Rs [14, 15]. RIC-3, a protein originally identified in the worm *Caenorhabditis elegans*, was shown to be necessary for proper functional expression of homomeric nAChR α7 and to enhance the surface expression of 5-HT_3A_R in HEK cells [16–18]. In contrast, co-expression of RIC-3 with heteromeric nAChR α3β4 and α4β2 as well as 5-HT_3A_R in *Xenopus laevis* oocytes inhibits the functional maturation of these receptors [15, 16, 18, 19]. The discrepancy in modulatory effects of RIC-3 on different pLGICs might stem from host cell variations and/or species differences of studied receptors [18, 20]. Previously, we have demonstrated that the intracellular domain of 5-HT_3A_ receptors (5-HT_3A_-ICD) is required for receptor modulation by RIC-3 because the removal of the ICD eliminated the inhibitory effects of RIC-3 co-expression on 5-HT_3A_ functional surface expression in *X. laevis* oocytes [21]. Further, we designed a chimera in which we added the 5-HT_3A_-ICD of 115 amino acids to the prokaryotic homologue of 5-HT_3A_, the *Gloeobacter violaceus* ligand-gated ion channel (GLIC), to substitute the receptor’s heptapeptide linker between M3 and M4 transmembrane helices. This addition lead to the observation of modulatory effects on expression of GLIC-5-HT_3A_-ICD chimera when co-expressed with RIC-3 while wild type GLIC was insensitive to co-expression [22, 23]. Additionally, we demonstrated that the interaction between GLIC-5-HT_3A_-ICD and RIC-3 is direct and does not require the presence of additional proteins. Both proteins were over-expressed in and purified from *Escherichia coli* (*E. coli*) and were subjected to a pull-down assay through which we observed an interaction between the two proteins [24]. Subsequently, in a separate study we examined if the 5-HT_3A_-ICD alone is sufficient to retain an interaction with RIC-3. We used a chimera obtained by fusing the 5-HT_3A_-ICD to the C-terminus of maltose binding protein (MBP). Addition of MBP to the 5-HT_3A_-ICD, facilitated the expression, purification and stability of this domain. We determined, using a similar pull down assay, that the purified 5-HT_3A_-ICD maintains an interaction with RIC-3 [25].

In summary, from previous studies, it is known that the 5-HT_3A_-ICD is required and sufficient for interaction of the 5-HT_3A_R and RIC-3. Furthermore, this direct interaction between the ICD and RIC-3 does not require the presence of other host cell proteins [21–25]. In the current study, we aimed to identify the segment of the ICD responsible for this interaction with the hypothesis that the interaction site lies within an independent segment of the ICD. In 2014, the X-ray structure of the mouse 5-HT_3A_R was published. The complete ECD and TMD were resolved, while part of the ICD (62 amino acids) was missing due to trypsin treatment of the receptor prior to crystallization. This unresolved segment of the ICD, here named L2, is located between the MX-helix that follows a short post M3 loop (L1), and the MA-helix that continues into M4 [26]. Recently, structures of full-length 5-HT_3A_R obtained by single-particle cryo-electron microscopy have been published, however, almost the entirety of the L2 segment remained unresolved [27, 28]. For the purpose of this study, we used the X-ray structure as a guide to design our constructs. Here, we utilized a process of elimination by dividing the ICD into its known structural elements (L1-MX, L2, and MA) (Figure 1), and determined whether RIC-3 interacted with the different segments. Through our experiments we demonstrate that the L1-MX region within the 5-HT_3A_-ICD interacts with RIC-3. Our results identified a stretch of 24 amino acids within the 115 amino acid long 5-HT_3A_-ICD as the interaction site of RIC-3.

**Figure 1.**
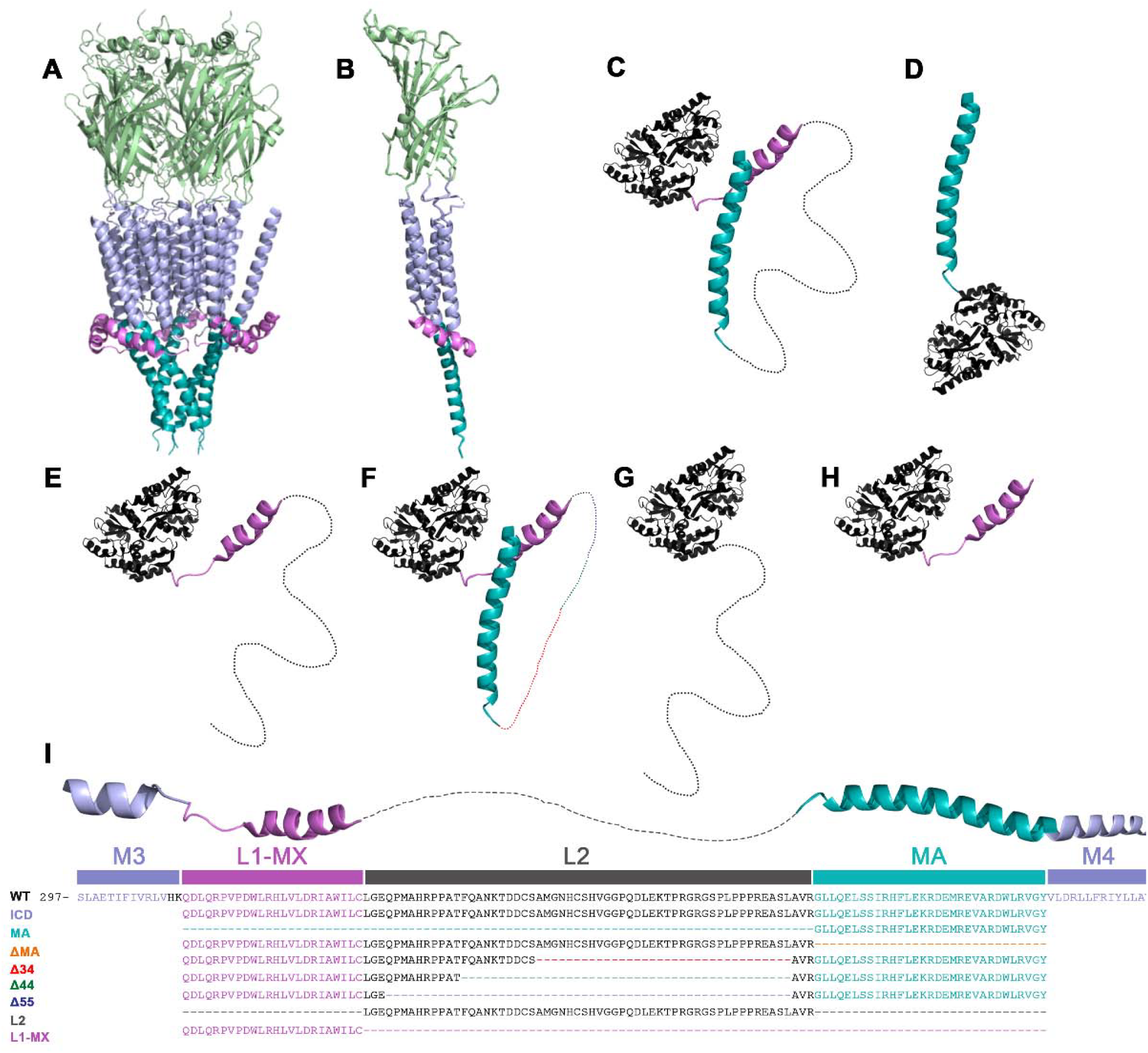
Structural representation and alignments of constructs. **A)** Cartoon representation of the pentameric m5-HT_3A_R X-ray structure viewed parallel to the plane of the membrane. ECD in green, TMD in purple, MA-helix in violet and MX-helix in cyan. **B)** A single subunit of the 5-HT_3A_ receptor viewed parallel to the membrane. **C)** Cartoon representation of MBP-5HT_3A_-ICD-WT. The cytoplasmic domain of 5-HT_3A_R is fused to the C-terminus of MBP, black structure, and serves as the template for respective constructs. The relative orientation of MBP with regard to the ICD is entirely hypothetical and only meant for overall illustration purposes. The 62 residues removed from L2 (charcoal) are shown as a dashed line. **D)** The MBP-MA has the N-terminus of the MA-helix attached to MBP. **E)** The MBP-ΔMA has the entire MA-helix removed. **F)** MBP-∆34, 44, 55 constructs are yielded by deleting 34 (red), 44 (green), and 55 (dark blue) amino acids from L2, respectively. **G)** The MBP-L2 contains only the unresolved segment of the ICD attached to the C-terminus of MBP. **H)** The MBP-L1-MX contains only the short loop and the helical structure post the M3 transmembrane segment. **I)** Multiple sequence alignment of 5-HT_3A_R and the ICD constructs highlighting deletions (dashed lines).

## Materials and Methods

### A. Materials

BL21-CodonPlus-(DE3)-RIPL cells (Agilent Technologies, Santa Clara, CA); ampicillin (Fisher Scientific, Fair Lawn, NJ); chloramphenicol (Fisher Scientific, Fair Lawn, NJ); IPTG (Fisher Scientific, Fair Lawn, NJ); leupeptin (AdipoGen Life Sciences, San Diego, CA); pepstatin (AdipoGen Life Sciences, San Diego, CA); PMSF (Research Products International, Mt. Prospect, IL); TCEP-HCl (Oakwood Chemical, N. Estill, SC); lysozyme (MP Biomedicals, Solon, OH); Protease inhibitor cocktail III (Research Products International, Mt. Prospect, IL); DNAse I (Alfa Aesar, Ward Hill, MA); DDM (Anatrace, Maumee, OH).

### B. Molecular biology

B.1. The cytoplasmic domain of the mouse 5-HT_3A_ receptor (QDLQ… RVGY, 115 amino acids) fused to the C-terminus of maltose binding protein (MBP) in pMALX vector (New England Biolabs), MBP-5-HT_3A_-ICD-pMALX, was used as the template to design constructs for the present study [25, 29]. The fusion constructs were generated by deletion of amino acids from the 5-HT_3A_-ICD using appropriate partially overlapping primers with the QuikChange II Site-Directed Mutagenesis kit (Agilent Technologies) and were confirmed by DNA sequencing (GENEWIZ, South Plainfield, NJ) [30].

B.2. Human RIC-3 (<u>NM_001206671.2</u>) in the pGEMH19 (hRIC-3-pHEMH19) expression vector was generously provided by Dr. Millet Treinin (Hebrew University, Israel). RIC-3 was cloned into the prokaryotic expression vector pMAL-c2x fused to N-terminal MBP and C-terminal His_6_ tag [24, 31, 32]. The MBP-RIC-3-His_6_-pMAL-c2x and hRIC-3-pHEMH19 were used in pull-down and *in-vivo* interaction assays, respectively.

B.3. For *in-vivo* interaction assays the 5-HT_3A_ subunit containing the V5 epitope tag (GKPIPNPLLGLDSTQ) close to the N-terminus, and the 5-HT_3B_ subunit were engineered into the expression vector pGEMHE and were used for full receptor expression in oocytes [21]. These cDNAs along with cDNA of hRIC-3-pHEMH19 were linearized with the restriction enzyme *Nhe*I (New England Biolabs). Subsequently, cRNA of each construct was prepared using T7 RNA polymerase (mMESSAGE mMACHINE^®^ T7 Kit; Applied Biosystems/Ambion, Austin, TX) in an *in-vitro* transcription process. The cRNAs were then purified and precipitated with MEGAclear^™^ kit (Applied Biosystems/Ambion, Austin, TX) and dissolved in nuclease-free water.

### C. Oocytes

Fresh oocytes used in *in-vivo* interaction assays were harvested from *Xenopus laevis* frogs and defolliculated in house. Maintenance and surgery procedures were approved by the TTUHSC animal welfare committee. Prior to injection with an automatic oocyte injector (Nanoject II™; Drummond Scientific Co., Broomall, PA) the oocytes were washed using Ringer’s buffer (OR2: 82.5 mM NaCl, 2 mM KCl, 1 mM MgCl2, 5 mM HEPES, pH 7.5). Injected oocytes were maintained in standard oocyte saline medium (SOS: 100 mM NaCl, 2 mM KCl, 1 mM MgCl2, 1.8 mM CaCl2, 5 mM HEPES, pH 7.5) supplemented with a 1% antibiotic-antimycotic (100x) and 5% horse serum at 15°C.

### D. Protein expression and purification

#### D.1. MBP-5-HT_3A_-ICD-pMALX fusion proteins

The MBP-5-HT_3A_-ICD-pMALX soluble fusion proteins were overexpressed in *Escherichia coli* (*E. coli*) BL21-CodonPlus (DE3)-RIPL cells (Agilent Technologies; Santa Clara, CA). The cells were harvested by centrifugation (5,000 g for 15 min at 4°C) and lysed in a two-step process using freshly prepared buffer (Buffer A: 20 mM Tris, 150 mM NaCl, 1 mM ethylenediaminetetraacetic acid (EDTA), 1 mM tris(2-carboxyethyl)phosphine (TCEP), pH 7.4). First the cell pellets were resuspended by stirring for 2 hrs at 4°C using buffer A containing added protease inhibitors (1 mM phenylmethylsulfonyl fluoride (PMSF); Research Products International, 10 μg/mL leupeptin; Sigma-Aldrich, 10 μg/mL pepstatin; Fisher Scientific), and 1 mg/ml lysozyme (Sigma Aldrich). The cell lysate solution was frozen at −80 °C overnight. Next, 50 μg/mL of DNaseI (Sigma Aldrich) and 1 mM PMSF were added to the thawed cell lysate solution and the mixture stirred for 2 hrs at 4°C. The cell suspension was passed through a 15 G needle several times to create a homogeneous solution. A clear soluble fraction of the lysate was obtained by centrifugation at 30,000 g for 1 hr at 4°C. The supernatant was passed through a 0.2 µm pore-size bottle-top filter and then loaded onto a gravity-packed amylose-resin column. The unbound proteins were washed extensively using buffer A (3 × 10 bed volumes) and the bound proteins were eluted by buffer A, pH 7.4, containing 20 mM maltose.

#### D.2. MBP-RIC-3-His_6_

The MBP-RIC-3-His_6_-pMAL-c2x membrane protein was overexpressed in *Escherichia coli* (*E. coli*), BL21-CodonPlus (DE3)-RIPL cells (Agilent Technologies; Santa Clara, CA). The cells were harvested by centrifugation (5,000 g for 15 min at 4°C) and lysed in a two-step process similar to above using freshly prepared buffer (Buffer A’: 20 mM Tris, 100 mM NaCl, 2 mM EDTA, 5 mM TCEP, 10% glycerol, pH 7.4). Following the lysis, the debris and unlysed cells were separated from the cell lysate solution by centrifugation (9,000 g for 20 min at 4°C). Next, cell membranes were collected by centrifugation of supernatant at a higher speed (30,000 g for 1 hr at 4°C). The membranes were re-suspended in solubilization buffer (Buffer B: 20 mM Tris, 100 mM NaCl, 5 mM TCEP, 10% glycerol, pH 7.4) containing 0.5% *n*-dodecyl β-D-maltoside (DDM). The solubilized cell membranes were further cleared by centrifugation (30,000 g for 1 hr at 4°C) and the supernatant was loaded onto a gravity-packed amylose-resin column. The resin was washed extensively with buffer B containing 0.05% DDM (3 × 10 bed volumes) and bound proteins were eluted with a solution of 20 mM maltose and 0.05% DDM in buffer B. Eluted proteins were subjected to a second step of purification by a gravity-packed Talon-resin column. The resin beads were equilibrated prior to adding the protein solution using cold buffer (Buffer C: 20 mM Tris, 500 mM NaCl, 5 mM TCEP, 10% glycerol, 0.05% DDM, 0.5% Triton x-100, pH 7.5) containing 10 mM imidazole. After 30 minutes incubation and gravity flow of the protein, the column was washed with Buffer C containing 15 mM imidazole. Next, the protein-bound resin bed was washed with a Triton x-100 free buffer (Buffer D: 20 mM tris, 500 mM NaCl, 5 mM TCEP, 10% glycerol, 0.05% DDM, pH 7.5) containing 15 mM imidazole. The bound protein was then eluted with 10 mL of the 400 mM imidazole solution in buffer D.

### E. Size exclusion chromatography

The amylose/Talon column purified proteins were subjected to size exclusion chromatography for final purification on a SuperdexTM 200 10/300 GL column. The column was equilibrated with SEC buffer (Buffer S: 20 mM Tris, 150 mM, NaCl, 1 mM TCEP, 5mM Maltose, 0.01% NaN3, pH 7.4 for MBP fusion constructs; Buffer S’: 20 mM tris, 500 mM NaCl, 5 mM TCEP, 10% glycerol, 0.05% DDM, 0.01% NaN3, pH 7.5 for MBP-RIC-3-His_6_). UV absorbance at 280 nm (*A*280) was utilized to monitor the elution of the proteins and fractions were analyzed after separation on 4– 15% precast gradient TGX Stain-Free^TM^ gels (Bio-Rad Laboratories).

### F. Pull-down assay

In this assay, 0.05% DDM and 1X protease inhibitor cocktail (Research Products International) were added to all buffers just before use in each step. First, 2.5 µg of freshly SEC-purified MBP-RIC-3-His_6_ (bait) was added to 15 µL of HisPur^TM^ Cobalt resin (Thermo Scientific) equilibrated with and re-suspended in 200 µL of binding buffer (Buffer A: 20 mM Tris, 150 mM NaCl, 1 mM MgCl2, 0.05% Triton X-100, 10 mM imidazole, pH 7.4) inside a Pierce™ Spin Cup - Cellulose Acetate Filter system (Thermo Scientific). After 30 minutes of incubation at 4°C, unbound MBP-RIC-3-His_6_ was washed away with 300 µL of wash buffer (Buffer B: 20 mM Tris, 150 mM NaCl, 1 mM MgCl2, 0.05% Triton X-100, 15 mM imidazole, pH 7.4) and centrifugation at 1,000 g for 30 seconds. This step was repeated five times and the resin re-suspended with 200 µL of buffer B. Next, the prey proteins (MBP-5-HT_3A_-ICD-pMALX constructs) were added to each individual spin cup 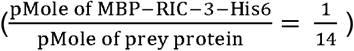 and mixtures were incubated at 4°C for 2 hours. Additionally, similar conditions were prepared using only the prey proteins with 15 µL of Cobalt resin to serve as controls. Unbound proteins were washed 12 times with 300 µL of buffer B and then eluted with 25 µL of elution buffer (Buffer C: 20 mM Tris, 150 mM NaCl, 1 mM MgCl2, 0.05% Triton X-100, 200mM imidazole, pH 7.4) and centrifugation at 1,200 g for 2 minutes. The eluted samples were immediately analyzed by SDS polyacrylamide gel electrophoresis (SDS-PAGE) using 4–15% precast gradient TGX Stain-Free^TM^ gels (Bio-Rad Laboratories, Inc.).

### G. In-vivo interaction assay

The *in-vivo* interaction assay was designed to probe the interaction of 5-HT_3A_ and 5-HT_3AB_ receptors with RIC-3. 1.5 ng of hRIC-3-pGEMH19 cRNA was co-injected with 6 ng of 5-HT_3A_-wt cRNA and 6 ng of 5-HT_3AB_-cRNA (3:1 ratio of A:B) into *Xenopus* oocytes, respectively. Oocytes injected with 5-HT_3A_-wt cRNA or 5-HT_3AB_ cRNA alone were used as controls. Current amplitudes induced by application of 10 µM of 5-HT in OR2 were measured 48 hours after the injection using Two-Electrode Voltage-Clamp (TEVC) recordings. Currents were recorded from individual oocytes perfused with OR2 at a membrane potential of −40 mV. They were amplified using a TEV-200A amplifier (Dagan Corporation; Minneapolis, MN), digitized using a Digidata^®^ 1440A analog-to-digital converter (Molecular Devices; Sunnyvale, CA), and analyzed with pCLAMP (Clampex/Clampfit) software (Molecular Devices).

### H. Statistics

Statistical significance was determined with one-way ANOVA and Tukey’s multiple comparisons test (**** denotes p < 0.001) of oocytes injected with 5-HT_3A_ or 5-HT_3AB_ and RIC-3 *vs* 5-HT_3A_ or 5-HT_3AB_ alone (Prism 6, GraphPad Software, Inc.).

### I. SDS-PAGE

For SDS-electrophoresis, 4–15% precast gradient TGX Stain-Free^TM^ gels (Bio-Rad Laboratories) were used and visualized by stain-free enabled imager (Gel Doc^TM^ EZ Imager, Bio-Rad).

## Results

In order to further investigate the interaction between the 5-HT_3A_R and RIC-3 and determine the interaction site within the 5-HT_3A_-ICD, we designed a series of complementary deletion constructs of the 5-HT_3A_-ICD in a stepwise manner (Figure 1). Initially, we engineered two deletion constructs based on structural elements observed within the 5-HT_3A_-ICD in the crystal structure of the mouse 5-HT_3A_ receptor [26]. We hypothesized that the interaction with RIC-3 involves specific and likely independent segments of the 5-HT_3A_-ICD. The crystal structure of the 5-HT_3A_ contains a partially resolved ICD with three distinct segments: the MX-helix following a short loop (L1) post the M3 segment of the TMD; the unresolved loop in between the two MX- and MA-helices (L2), and the MA helix, the 31 amino acids long helical structure preceding the M4 segment of the TMD. To obtain the first construct we deleted the MA-helix from the 5-HT_3A_-ICD, yielding ΔMA (QDLQ…LAVR). The second construct was obtained by deletion of L1-MX-L2 leaving only the MA-helix, MA (GLLQ…RVGY). Three additional deletion constructs were designed comparable to a previous study investigating the functional impact of deletions of the ICD [33]: Δ34 was generated by deletion of 34 amino acids from the C-terminus of L2 (∆A358-L392); Δ44 was generated by deletion of 44 amino acids from the C-terminus of L2 (∆F348-L392); Δ55 was generated by deletion of 55 amino acids from the C-terminus of L2 (∆E337-L392); L1-MX was generated by deletion of L2 and MA (QDLQ…WILC); and L2 was generated by deletion of L1-MX and MA (LGEQ…LAVR). All constructs were fused to MBP at their N-terminus forming soluble constructs. Next, we over-expressed the constructs in *Escherichia coli* (*E. coli*) and purified the cell lysates utilizing an amylose resin affinity column for the first step of purification. The eluted fractions were then analyzed using SDS-PAGE (Figure 2A). The protein solution was then subjected to size exclusion chromatography (SEC) for further purification, and purified protein fractions were visualized by SDS-PAGE (Figure 2B). The elution volumes of different ICD constructs upon SEC indicate different oligomeric states of different ICD constructs (Figure 2C and 2D). We have characterized and identified a potential oligomerization motif related to the conductance-limiting arginines within the MA-helix and an aspartate within L1 (bioRxiv, doi: https://doi.org/10.1101/561282, under review). It is important to mention that subsequent deletions of amino acids in designing further constructs did not cause any significant changes in pI of the proteins, and therefore, the same expression and purification conditions were used for all MBP-5-HT_3A_-ICD constructs.

**Figure 2.**
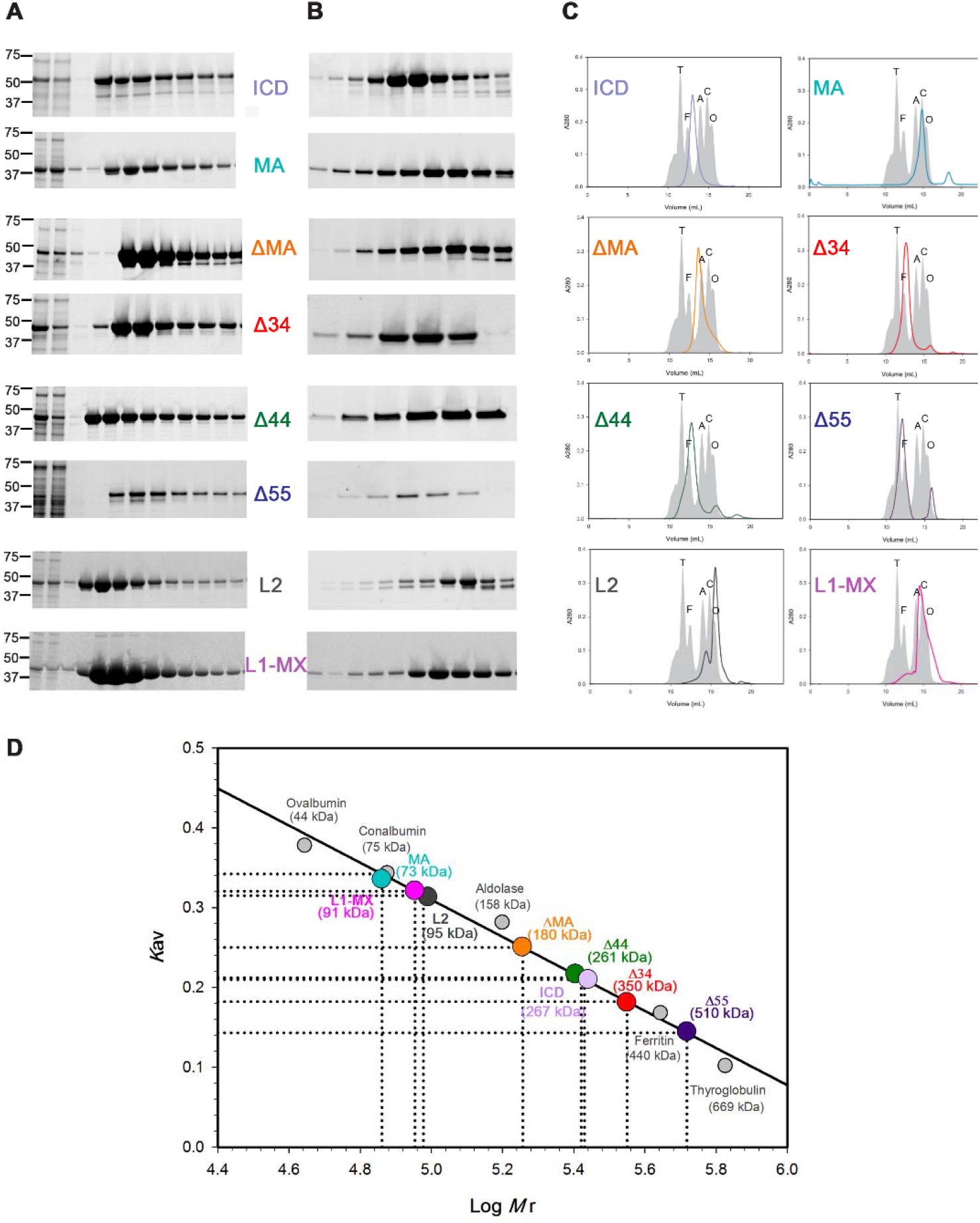
Two step purification of the ICD constructs. **A)** Proteins were purified by amylose-resin column chromatography. The soluble protein was loaded onto columns (1^st^ lane: loading material), and was passed through via gravity flow (2^nd^ lane: flow-through), followed by washes (3^rd^ lane)), and eluted in fractions (remaining lanes). S tain-free SDSTGX-gels indicate the quality of purified protein, identify peak fractions and determine the weight of denatured monomeric ICD constructs. From top: ICD (53 kDa), MA (45 kDa), ΔMA (49 kDa), Δ34 (50 KDa), Δ44 (48 kDa), Δ55 (46 kDa), L2 (45 KDa), and L1-MX (42 KDa). A standard protein ladder is shown next to each protein. B) Stain-free SDS-TGX-gel of peak fractions of each construct after size exclusion chromatography (SEC). **C)** SEC Chromatograms for each construct against the standard proteins, grey area, in their respective colors. **D)** Standard curve used for determination of molecular weight and oligomeric state of each construct. Grey circles represent standard proteins, and constructs

We additionally overexpressed the MBP-hRIC-3-His_6_ chimera [24] in *E. coli* and optimized the lysis and purification of this membrane protein to obtain a homogenous and stable protein with appropriate yield for our direct protein-protein interaction studies. RIC-3 was purified using two sequential affinity purification steps with amylose and Talon super-flow resins and additionally SEC. The solubilized protein is observed at 100 kDa on SDS-PAGE electrophoresis (theoretical Mw: 84.9 kDa) (Figure 3)

**Figure 3.**
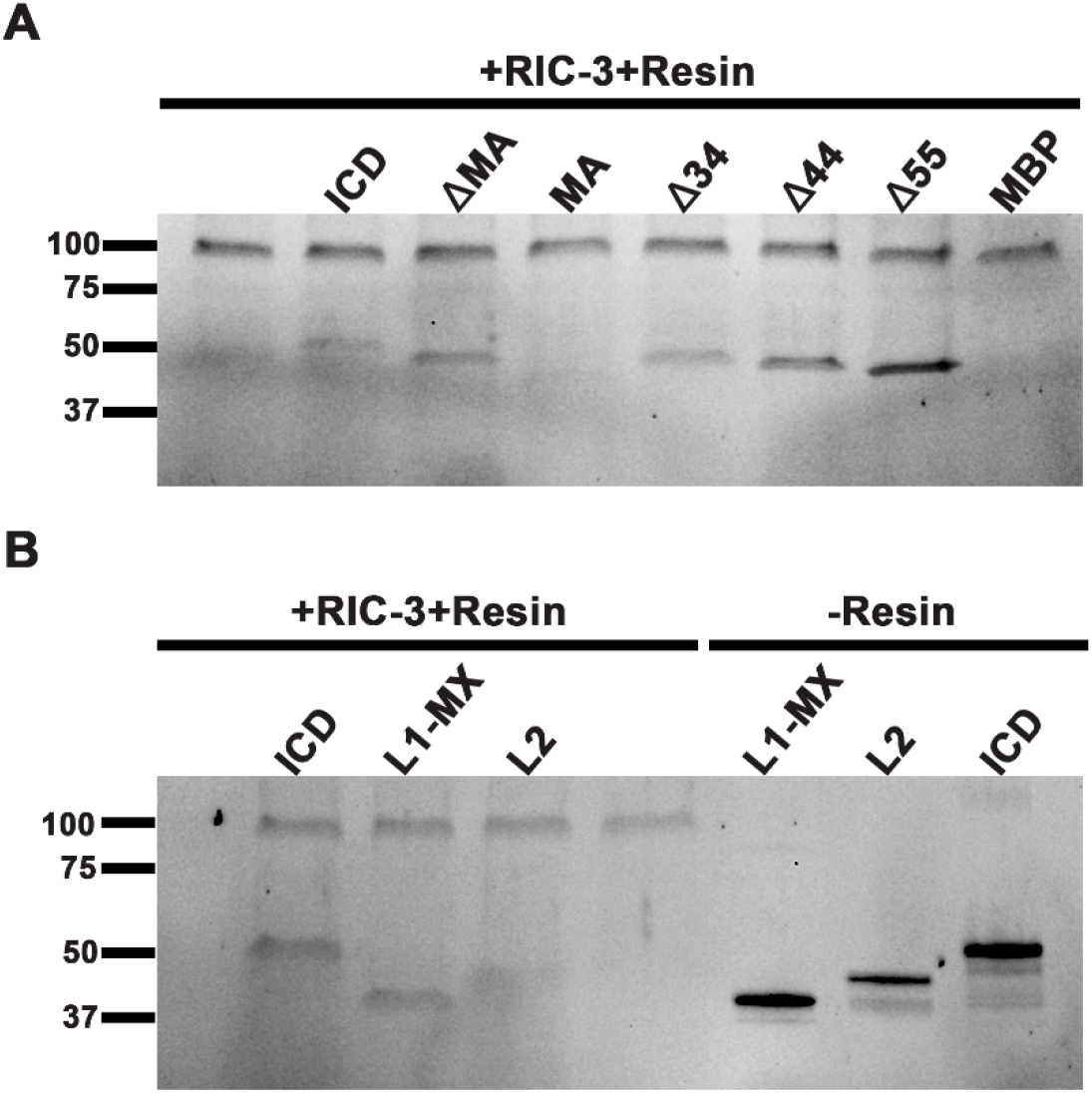
RIC-3 pull-down assay. To assess protein-protein interaction each 5-HT_3A_-ICD construct (prey) was incubated with purified MBP-RIC-3-His_6_ (bait) bound to Cobalt resin. Complexes retained on beads were eluted and analyzed using Stain-free SDS-TGX gel electrophoresis. MBP-RIC-3-His_6_ is observed at 100 kD (upper band) and the presence of an interaction is indicated by appearance of a second band corresponding to the size of the protein of the interest. Lack of a lower band indicates no interaction with RIC-3. Experiments were repeated several times and we observed the same results. **A)** SDS-PAGE analysis of eluted complexes as follows: Lane 1, RIC-3 only to ensure binding to the resin; lane 2-8, eluates from pull-downs with RIC-3 and constructs as indicated. **B)** SDS-PAGE analysis of eluted complexes as follows: Lane 1-3, eluates from pull-downs with RIC-3 and constructs as indicated; Lane 4, RIC-3 only to ensure binding to the resin. The last 3 lanes are control proteins to indicate the localization on SDS-PAGE without either resin or RIC-3.

### Effect of deletions on interaction with RIC-3

We investigated the interaction between the ICD constructs and RIC-3 by conducting pull-down assays directed against the Histidine tag of the RIC-3 construct. The resin bound RIC-3 protein was incubated with equal molar amounts of ICD, ΔMA and MA constructs, respectively, and the eluates were analyzed by SDS-PAGE under reducing conditions to visualize both the bait and prey proteins (Figure 3). Under these conditions, observing only one single band on the gel corresponding to the size of RIC-3 (bait) migrating as a major band with a relative molecular weight of 100 kDa indicates that there was no interaction between RIC-3 and the added ICD construct. On the other hand, observing two bands, one corresponding to the size of RIC-3 and one to the size of the respective chimera illustrates the presence of both bait and prey protein and therefore an interaction. We observed a second band in the SDS-PAGE gel in conditions where RIC-3 was incubated with the ICD (~53 KDa) or ΔMA (~50 KDa), indicating interactions of the involved proteins. On the contrary, no significant band was observed for the MA construct (~41 KDa) (Figure 3A). The direct interaction of the full-length ICD and RIC-3 had been shown previously in a similar experiment and the ICD construct here was used as a control [25]. MBP alone or omitting the RIC-3 in the pull-down did not reveal the presence of the prey protein.

Eliminating the MA-helix from potential interaction sites of the ICD with RIC-3 shifted our focus to ΔMA which contains L1, MX, and the structurally unresolved L2 segments. We designed three constructs to understand the role of L2, the disordered flexible region between MA- and MX-helices, in the observed interaction between RIC-3 and ΔMA: Δ34, Δ44, and Δ55. Using the pull-down assay, we observed strong interactions between all three deletion constructs and RIC-3 using SDS-PAGE analysis (Figure 3A). The second band for each condition was observed at relative molecular weights of 50 kDa for Δ34, 48 kDa for Δ44, 46 kDa for Δ55, consistent with the calculated weights of the prey proteins (Table 1). Of note, we observed an increase in intensity of these prey bands as the number of deleted amino acids increased. The L2 region of the ICD is formed by 60 amino acids, and by deleting 55 amino acids in Δ55 construct, we eliminated most of this region from the ICD. Therefore, we hypothesized that the interaction site must lie within the remaining segment of ΔMA; the L1-MX region.

**Table 1:** Constructs and respective molecular weight and number of amino acids

| Construct | Theoretical molecular weight of<br>the monomer fused to MBP<br>(M<sub>w</sub>, 10<sup>3</sup> g/mole) | Number of<br>amino acids |
| --- | --- | --- |
| ICD | 53.4 | 115 |
| MA | 44.3 | 31 |
| ΔMA | 49.8 | 84 |
| $\Delta 34$ | 50.0 | 81 |
| $\Delta 44$ | 48.9 | 71 |
| $\Delta 55$ | 47.7 | 60 |
| L2 | 46.8 | 60 |
| L1-MX | 43.4 | 24 |

### Identifying the short peptide responsible for RIC-3 interaction

We continued our studies following our initial strategy of stepwise complimentary deletion of amino acids. Our final constructs where designed by using ΔMA as the template. We deleted the L2 region from ΔMA to create the L1-MX construct and, to complement this construct, we deleted the L1-MX segment to create the L2 construct. These constructs were designed to confirm that the L2 region does not confer the interaction of 5-HT_3A_ and RIC-3 and that L1-MX is independently responsible for mediating this interaction. We performed the RIC-3 pull-down assay and observed a second band corresponding to the size of L1-MX (42 kDa) and no significant band in case of the L2 construct (Figure 3B).

### *In-vivo* assay comparing interaction of homomeric and heteromeric 5-HT_3_ receptors with RIC-3

A sequential alignment between 5-HT_3A_ and 5-HT_3B_ subunits of 5-HT_3_ receptors revealed that 5-HT_3B_ potentially lacks the helical structure post M3, the MX-helix (Figure 4A). Since we had just demonstrated that the MX-helix mediates the RIC-3 modulation of 5-HT_3A_ subunits, we wanted to determine whether RIC-3 co-expression impacts functional heteromeric 5-HT3_AB_ expression similar to homomeric 5-HT_3A_ expression. 5-HT_3B_ subunits alone are unable to form functional channels. Co-expression of 5-HT_3A_ with 5-HT_3B_ is required for functional expression [34]. We used the cRNA of 5-HT_3A_ and 5-HT_3B_ subunits to express homomeric 5-HT_3A_ and heteromeric 5-HT_3AB_ in *Xenopus* oocytes. Both 5-HT_3A_ and 5-HT_3AB_ receptors were also co-expressed with human RIC-3 (hRIC-3) in oocytes. The current amplitudes of the receptors in each condition were measured in response to 10 µM 5-HT by two electrode voltage clamp technique (TEVC) 48 hours post injection (n ≥ 3) (Figure 4B and C). There was no significant difference (P = 0.93) in Mean + SEM of currents measured from oocytes injected with 5-HT_3A_ vs 5-HT_3AB_. We observed a significant drop in current amplitudes in oocytes co-injected with 5-HT_3A_ and hRIC-3 (P ≤ 0.001) as well as in oocytes co-injected with 5-HT_3AB_ and hRIC-3 (P ≤ 0.001) compared to the respective 5-HT3 injected alone. The difference in inhibition in the latter two conditions were not statistically significant (P ≤ 0.001).

**Figure 4.**
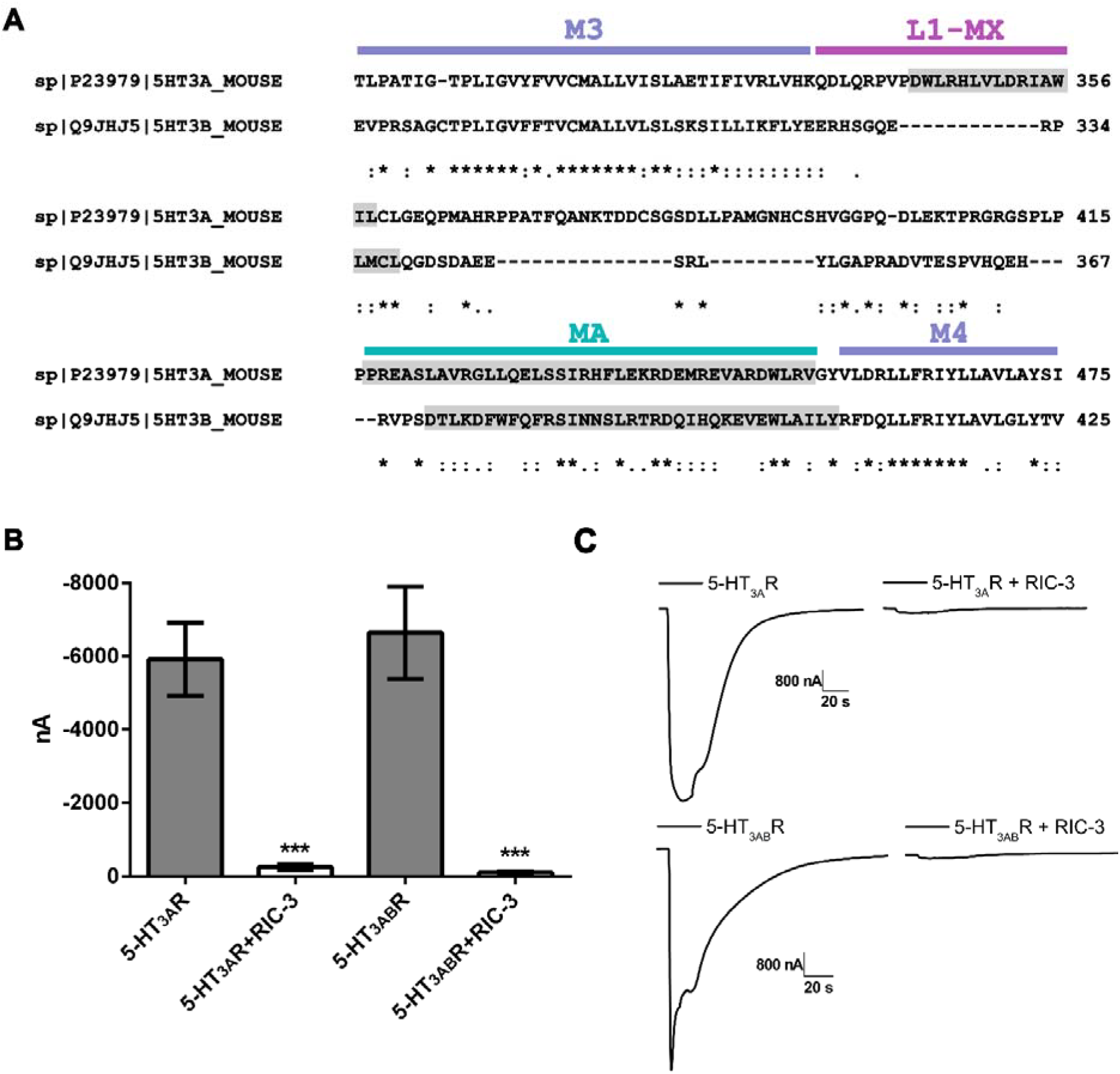
Comparison of homomeric and heteromeric 5-HT_3_Rs in RIC-3 interaction. **A)** Secondary structure prediction of 5-HT_3A_ and 5-HT_3B_ subunits obtained with PSIPRED. Predicted α-helical segments of the ICDs are highlighted in grey. 5-HT_3A_ contains a 15-amino acid long helical segment post M3 whereas the 5-HT_3B_ subunit only has 4 helical amino acids. **B)** 10 µM 5-HT induced current amplitudes (nA) were measured 48 hours after injection of cRNA into X. laevis oocytes by two-electrode voltage-clamp experiments. Currents recorded for oocytes injected with 5-HT_3A_ and 5-HT_3AB_ (grey bars) and after co-expression together with hRIC-3 (white bars). Statistical significance was determined with one-way ANOVA and Tukey’s multiple comparison test (*** denotes p < 0.001). n>3 oocytes were used for each condition. Currents were almost completely abolished after co-expression with RIC-3 for both 5-HT_3A_ and 5-HT_3AB_. **C)** Sample traces of currents for each condition are representative of data used for panel B.

## Discussion

With more than 40 different subunits of the pLGIC super family found in humans, there is a great abundance of these ion channels in neuronal tissues and some expression in non-neuronal cells and tissues [13]. The role of pLGICs is extremely important in signaling mechanisms and functioning of the nervous system, and disruptions to their function lead to a wide variety of diseases and disorders [9–11, 13]. According to existing structures of several pLGICs, such as 5-HT_3_, several nAChRs, GABA_A_β3 and Glyα1 receptors, there is a high conservation in composition and structure of the ECD and TMD of these receptors **[26–28, 35–38]**. The commonality of most of these structures is that intracellular domains (ICD) have been removed, shortened, or in case of full-length receptor structures, the ICD remained partially resolved [26, 27]. Additionally, all current drugs targeting these receptors interact with either the ECDs or TMDs **[5, 7–9, 39]**. Considering the significant sequence similarities of these domains among the super family, drugs interacting with these two conserved domains have been shown to cause undesired side effects by interacting with related subunits. Interestingly, ICDs of these receptors are divers in length, 73 to 262 amino acids in humans, and amino acid composition. This indicates that targeting the ICD may yield subtype-specific drugs. Therefore, the ICDs may represent safer and more effective potential drug targets for many diseases and disorders **[8, 40]**. Resistance to inhibitors of cholinesterase (RIC-3) is a chaperon protein known to modulate maturation and membrane trafficking of some members of pLGICs. Previously we have shown that the ICD mediates this interaction [21–23], and that the ICD alone, in the absence of both ECD and TMD, interacts with RIC-3[25].

In the present study, we aimed to identify the segment of the ICD involved in the interaction with RIC-3 using a series of constructs that were designed with different deletions of segments of the ICD. After the α-helical M3 transmembrane segment the ICD begins with a short loop (L1) that leads into an α-helical segment of about 18 amino acids in length (MX-helix), which is followed by a loop (L2) of 60 amino acids that has not been resolved in the structures. At the C-terminus of the ICD an α-helical segment of 31 amino acids (MA-helix) is continuous with the last transmembrane segment, M4. We had initially hypothesized that the MA-helix that is only present in cation-conducting pLGICs mediates the interaction with RIC-3. Using a direct protein-protein interaction assay we observed that RIC-3 directly interacts with ΔMA and not with the MA-helix segment of the ICD. These results indicated that the area of interaction lies within the L1-MX-L2-segment. In order to dissect a potential interaction site within the long L2 region, we designed three constructs that lacked 34, 44, and 55 amino acids, respectively, of L2 within the parent L1-MX-L2-MA construct. These three ICD constructs maintained interactions with RIC-3. Furthermore, a direct correlation between intensity of the interaction (observed by intensity of the corresponding band on the gel) and the number of amino acids deleted from the L2 region was observed. We suspect that the deletion of amino acids from the L2 region caused conformational changes that facilitated interaction between ICD and RIC-3. Consequently, we inferred that neither the MA-helix nor L2 are the primary contributor to interaction sites of the ICD with RIC-3. At this point we hypothesized that the L1-MX region contains the interacting segment. From the crystal structure of the 5-HT_3A_ it is possible to infer that the MX-helix would be responsible for mediating the interaction between the 5-HT_3A_ receptor and a membrane protein such as RIC-3. The conformation of the MX-helix in the structure and positional proximity of this helix to the membrane spanning helices creates a favorable position for protein-protein interaction. One caveat was mentioned in the paper: “However, whether the conformation of the post-M3 loop and of the MX helix in the detergent-solubilized, trypsin-treated, crystallized receptors do accurately represent a physiological conformation of intact receptors at the plasma membrane remains to be investigated”[26]. To assess our hypothesis, we designed two constructs, L1-MX and L2. The observed interaction between the L1-MX but not the L2 construct and RIC-3 indicated our hypothesis to be true. In conclusion, we have identified the 24 amino acid long L1-MX segment of the ICD to be the interaction site of the 5-HT_3A_ receptor and RIC-3.

As mentioned previously, chaperon protein RIC-3 has been shown to be a modulator of functional surface expression of 5-HT_3A_Rs, the only homomeric receptors in 5-HT_3_Rs family. However, since the 5-HT_3A_ subunit is required for assembly with other 5-HT(B-E) subunits to form heteromeric receptors, RIC-3 may have an effect on heteromeric formation of these receptors **[3, 13, 34, 41]**. In 2007 it was discovered that a specific isoform of RIC-3 enhances the surface expression of homomeric 5-HT_3A_Rs while inhibiting the expression of heteromeric 5-HT_3AB_Rs in COS-7 cells **[42]**. Shortly afterwards a separate study revealed that RIC-3 interacts with A, C, D and E 5-HT_3_ subunits and enhances the surface expression of 5-HT_3A_Rs in HEK cells **[43]**. These studies suggest that RIC-3 plays a role in the composition of 5-HT_3_Rs and is involved in folding, assembly, or transport of homomeric 5-HT_3A_Rs at the expense of heteromeric receptor formation **[18, 43]**. However, 5-HT3_AB_Rs are the only identified heteromeric receptors that resemble characteristics of native 5-HT_3_Rs in HEK cells **[34]**. In this study, we have identified the L1-MX segment of the 5-HT_3A_ subunit to be responsible for interaction between 5-HT_3A_Rs and RIC-3. Interestingly, according to a canonical sequence alignment between 5-HT_3A_ and 5-HT_3B_ subunits and secondary structure predictions using *PSIPRED* the 5-HT_3B_ lacks the MX-helix (Figure 5). Here, we investigated the effect of RIC-3 on heteromeric 5-HT3_AB_Rs to widen our understanding of these highly native-like receptors. We co-expressed the mRNA of both 5-HT3_A_ and 5-HT3_AB_ with hRIC-3 in *X*. oocytes and recorded the currents in response to 5-HT using TEVC. Co-expression of RIC-3 attenuated 5-HT-induced currents in both 5-HT_3A_ and 5-HT_3AB_ expressing oocytes, with no significant difference between 5-HT_3A_Rs and 5-HT_3AB_Rs. It has been suggested that heteromerization of 5-HT_3B_ subunits with 5-HT_3A_ subunits covers ER-retention signals found in 5-HT_3B_ subunits [44]. Our results, although not conclusive, suggest different possibilities leading to hRIC-3 attenuating surface expression of 5-HT3_AB_: RIC-3 may bind to unassembled 5-HT_3A_ subunits and inhibit their assembly with 5-HT_3A_ or 5-HT_3B_ subunits and thus inhibit both homomer and heteromer formation, or the interaction may block trafficking to the membrane.

Several lines of evidence, including a point mutation in the 5-HT_3A_-ICD, support the involvement of 5-HT_3_Rs in bipolar disorder and schizophrenia **[5, 9, 11, 12, 40]**. Interestingly, an increase in the level of RIC-3 mRNA has been shown in post-mortem brains of individuals suffering from these neurological diseases **[45]**. In the present study, we used a series of deletions and a direct protein-protein interaction assay to more closely define the interaction site of RIC-3 within the 5-HT_3A_-ICD. We identified a 24 amino acid long peptide, L1-MX, to bear the interaction site. Further studies are needed to identify the face of the α-helix or the exact amino acids responsible for this interaction and additionally to delineate the binding site on RIC-3. Our data provides a potential surface area for future drug design to specifically target 5-HT_3_ receptors and creates the basis for novel approaches for the treatment of these diseases.

## Acknowledgments

Research reported in this publication was supported by the National Institute of Neurological Disorders and Stroke of the National Institutes of Health under award number R01NS077114 (to MJ). The content is solely the responsibility of the authors and does not necessarily represent the official views of the National Institutes of Health. We thank the TTUHSC Core Facilities: some of the images and or data were generated in the Image Analysis Core Facility & Molecular Biology Core Facility supported by TTUHSC.

The authors declare no conflict of interest.

## AUTHOR CONTRIBUTIONS

M.J. designed research. E.P., A.G.S. and M.J. conceived the wild-type and engineered ICD constructs. E.P., A.G.S., A.P. and M.J. performed experiments and analyzed data. E.P., A.G.S., A.P., and M.J. wrote the paper.

